# Deep Learning-Based Motif Discovery in Major Histocompatibility Complex: A Primer

**DOI:** 10.1101/2024.10.03.616544

**Authors:** Bruno Alvarez, Morten Nielsen

## Abstract

In this work, we present a deep learning-based approach for motif discovery in the Major Histocompatibility Complex (MHC) system, which plays a key role in the immune response. We explore the use of convolutional neural networks (CNNs) to identify peptide binding motifs for different MHC alleles. By training models on data from specific MHC Class I and II molecules, we demonstrate how 1-dimensional convolutional filters can effectively capture motifs and binding preferences. Our study introduces a method for extracting motif logos directly from the trained models, providing insights into how internal neural network representations align with known biological motifs. The results show significant alignment with experimental binding motifs, underscoring the utility of deep learning in immunological research and the potential for improving vaccine design and immunotherapy.

## Introduction

The Major Histocompatibility Complex (MHC) is a foundational protein of the adaptive immune system that mediates the presentation of antigenic peptides to T cells. This process allows the immune system to detect and respond to a variety of pathogens and cellular abnormalities [Janeway et al., 2001]. MHC Class I (MHC-I) molecules, found on nearly all nucleated cells, present peptides derived from intracellular sources, while MHC Class II (MHC-II) molecules, primarily expressed by specialized antigen-presenting cells such as dendritic cells and macrophages, present peptides originating from extracellular sources [Neefjes et al., 2011]. Both MHC-I and MHC-II display these antigenic peptides on the cell surface to facilitate T cell recognition. MHC-I specifically presents these peptides to CD8+ cytotoxic T cells, leading to the elimination of infected or cancerous cells, whereas MHC-II presents peptides to CD4+ helper T cells, which play a crucial role in orchestrating broader immune responses, including antibody production and activation of macrophages.

The ability to determine which peptides bind to specific MHC molecules is critical for understanding immune recognition, predicting immune responses, and developing immunotherapies and vaccines. However, the highly polymorphic nature of MHC molecules, which enables the presentation of a diverse range of peptides, presents a significant challenge for experimental characterization. Methods such as X-ray crystallography and mass spectrometry, though informative, are resource-intensive and not feasible for large-scale analysis across the wide variety of MHC alleles [Sette and Rappuoli, 2008]. Early computational methods, such as Position-Specific Scoring Matrices (PSSMs) and sequence motif-based approaches, provided initial frameworks for predicting peptide-MHC interactions but had limited flexibility and accuracy due to their reliance on linear scoring and lack of consideration for complex binding features [Bui et al., 2005, Parker et al., 1994].

While early methods like PSSMs aimed to describe MHC binding in a straightforward, linear manner, modern machine learning techniques, such as Feed-Forward Neural Networks (FFNNs), have significantly improved these capabilities. Models like NNAlign-2.1 [Andreatta and Nielsen, 2016], which are trained on Single-Allelic (SA) datasets containing data for a single MHC allele, and NNAlign MA [Alvarez et al., 2019], trained on both SA and Multi-Allelic (MA) datasets that incorporate data from multiple MHC alleles, now excel at learning the complex, non-linear interactions between peptides and MHC molecules. In both cases, the FFNN architecture behind these algorithms effectively captures significant patterns from the input data, embedding this information within the network’s structure. Consequently, after training the network, it becomes feasible to generate MHC binding motifs by predicting a large number of random peptides (typically around 200,000), selecting the highest-scoring ones (usually the top 0.1-1%), and constructing a sequence logo to visualize these motifs. This approach is routinely applied, for example, to illustrate receptor preferences in the NetMHCpan-4.1 and NetMHCIIpan-4.0 motif viewers [Nielsen and Andreatta, 2020b,a]. It is important to note that these models often derive binding motifs post hoc, by analyzing and aligning the highest-scoring peptides only after the model has completed its training. Although effective, this workflow is still highly dependent on specific hyperparameters, such as P1 preference priming, which can limit the overall transparency of the model and hinder the direct interpretability of the learned internal representations

To illustrate how the approach of generating binding motifs has extended beyond the MHC domain, Fenoy et al. [2019] applied a similar technique to create kinase phosphorylation motif logos. These motifs were generated after constructing the NetPhosPan algorithm, a generic deep convolutional neural network designed for predicting ligand-receptor interactions. NetPhosPan was trained as a pan-specific peptide-kinase binding predictor, in much the same way that NNAlign MA was employed to predict pan-specific peptide-MHC interactions. The primary architectural difference between NNAlign MA and NetPhosPan lies in the implementation of 1-dimensional convolutional layers (CLs) in the latter, which are used to “scan” input kinase domains for potential binding sites. In brief, encoded kinase sequences are fed into a convolution module consisting of three parallel CLs with lengths (3, 5, 7) and 40 filters each. The global max-pooled outputs of these layers are then concatenated with the encoded ligand sequences and subsequently fed into a shallow FFNN with a single output neuron that predicts the phosphorylation likelihood for the submitted ligand-kinase pairs.

Due to its deep learning nature, the above example represents an intriguing case for further analysis. The rationale behind this is that convolutional layers possess the ability to slide over a sequence (which may be temporal, semantic, etc.) and adaptively “pay attention” to different regions across the input series (in the work of Fenoy et al., these regions correspond to the kinase binding domains). For this adaptive process to occur, the convolutional filter weights-characterized by matrices-are iteratively optimized using computational methods such as Gradient Descent. This optimization dynamically shapes the filter space, allowing the matrices to capture well-defined preferences for specific amino acid compositions at precise positions. Consequently, given such an enriched representation space, an interesting question arises: could receptor motifs be directly extracted from this space? This would entail constructing receptor binding motif logos directly from the network weights, rather than relying on the top predictions of random peptides as previously described.

To start exploring possible answers for the above question, a careful dissection and understanding of the 1D convolution operator must be achieved first. From a deep learning perspective, 1D convolutions are, essentially, sliding matrices that are applied to some input data matrix in order to detect repeating patterns. Formally, we can define such operation as [PyTorch, 2024]:

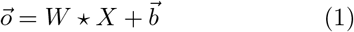

where 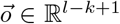 is the output vector, *W* ∈ ℝ^*c×k*^ is the weight matrix (or filter of kernel size *k*) of the convolution, ⋆ is the convolution operator, *X* ∈ ℝ^*c×l*^ is the encoding of some input sequence *s* (i.e. with BLOSUM) of length *l* (generated from an alphabet *A* of length *c*), and 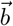 is the bias vector (refer to Figure 1 for a graphical example). With this, the ⋆ operator is defined to 1) compute the pairwise multiplication of all elements in *W* and *X* at a given sliding position, 2) sum all these multiplications (condensing everything into a single scalar), and 3) store this scalar in an output vector at the same index of the current sliding position.

**Figure 1:**
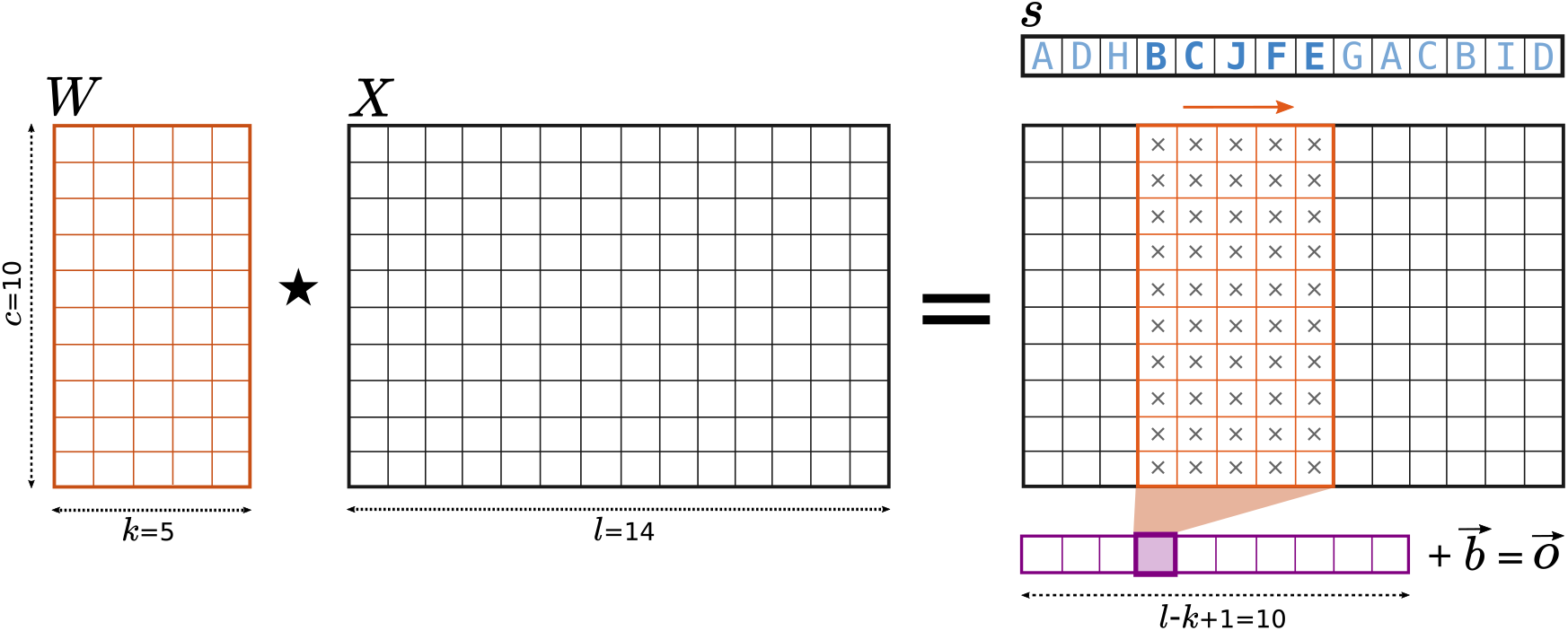
Toy example of a 1-dimensional convolution. Here, a filter *W* with a kernel size *k* of 5 and 10 channels *c* is being convolved with the encoding *X* of the hypothetical sequence *s* = ADHBCJFEGACBID, of length *l* = 14, and constructed via sampling of an alphabet of length *c. W* is slided from left to right over the position axis, and for each possible sliding step, the convolution ⋆ (the sum of all pairwise multiplications ×) is computed and stored in a vector of maximum length *l* − *k* + 1 = 10 (in purple). Then, this convolution vector is added to the bias 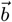, resulting in the final output 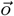. In this example, the filter is shown located “on top” of *X* at sliding position 3, with the convolution result stored in the position 3 of 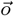.

As a result, the elements of 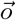 will carry information regarding the presence or absence of particular encoded characters *a* ∈ *A* at specific positions of *s*. Moreover, filters of different *k* values will scan *s* for specific subsequences of length *k*, making 1D convolutions a fit candidate for multi-resolution motif recognition. A rightful question, however, might be raised: why is that ⋆ is able to capture such information? The short answer is that, in essence, the 1D convolution is a sliding dot product operator. This means that a filter matrix *W* convolving *X* computes column-wise dot products in each sliding position, which span an intermediate vector 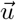, whose position-wise sum yields the final convolution output for such a position (see Figure 2 for a visual example). Since the dot product between two vectors is a way of measuring the projection of one onto the other, these column-wise dot products represent an explicit way of projecting elements of *s* onto a filter’s space. With this, 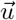 can be directly interpreted as the projection of sequence *s* onto a filter *f* (or, analogously, the projection of *X* onto *W*), at a specific sliding position. Since a *W* of size *k* can be placed at *l* − *k* + 1 different positions over *X*, a convolution will span a total of *l* − *k* + 1 possible output values. Among these values, there will be a maximum one, and it will be associated to a specific sliding position *p*. This position is of great importance, since it conveys the starting position of the subsequence of *s* which generated the greatest filter response.

**Figure 2:**
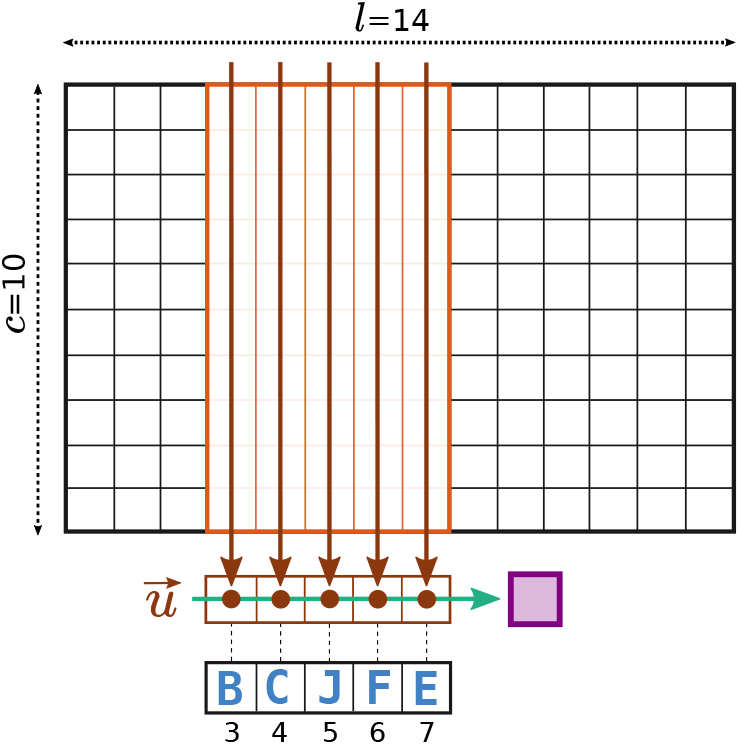
Computing a 1D convolution from a sliding dot product viewpoint. Here, the same situation from Figure 1 is shown. The convolutional filter *W* (in orange) is sitting on top of *X* (in black) at position 3, and extends itself up to position 7. In a column-wise manner, dot products between *W* and *X* are calculated and stored in an intermediate projection vector 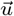 (in brown). Then, by summing the components of 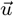 (green arrow) the convolution value for position 3 is obtained (purple block). Notice how there is a unique correspondence between elements of 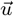, letters of the subsequence BCJFE, and the positions of such letters in *s*. A side note: another possible approach to compute the same convolution value is to do the dot product row-wise, and then summing (this, however, is not going to be covered here).

With this last piece of information, we can now associate this subsequence to a filter’s peak activation position *p*, and as a result recover the best projection vector as:

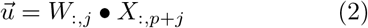

with 0 ≤ *j* ≤ *k* − 1 and • being the dot product. Notice how *j* indexes only column positions, meaning that the operation is conducted across rows (in the same way as displayed in Figure 2). If we now take into account that 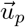 comes from a subsequence of *s*, we can remap the elements of this vector into a “projection matrix” whose rows are indexed by alphabet elements *a*, and columns by kernel positions.

Moreover, considering an input space *S* (with multiple sequences *s*), summing the projection matrices of each *s* will serve as a way of sampling a target convolution’s projection image. we will refer to this sampled image simply as projection, and denote it with *ϕ* according the following expression:

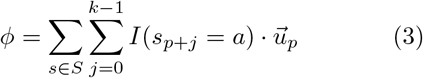

where *a* represents any symbol in alphabet *A* (in our case, the 20-letter amino acid alphabet), *s*_*p*+*j*_ is the symbol found at position *p* + *j* of *s*, and *I* is an indicator function [Wikipedia, 2024] of *s* (defined as 1 if *s*_*p*+*j*_ = *a*, 0 otherwise) used for indexing amino acids in *ϕ* (refer to Figure 3 for an example computation). Given the fact that a filter *W* will have a preference for specific amino acids at certain positions, *ϕ* will capture the presence/absence of such a preference in *S*, and, furthermore, give an estimate of its amplitude (by means of accumulation by summing for all *s*). Also, thanks to the indexing provided by *I, ϕ* will have akin characteristics to a PSSM, and thus it will be possible to treat it as a such (i.e. to generate a logo for visualization).

**Figure 3:**
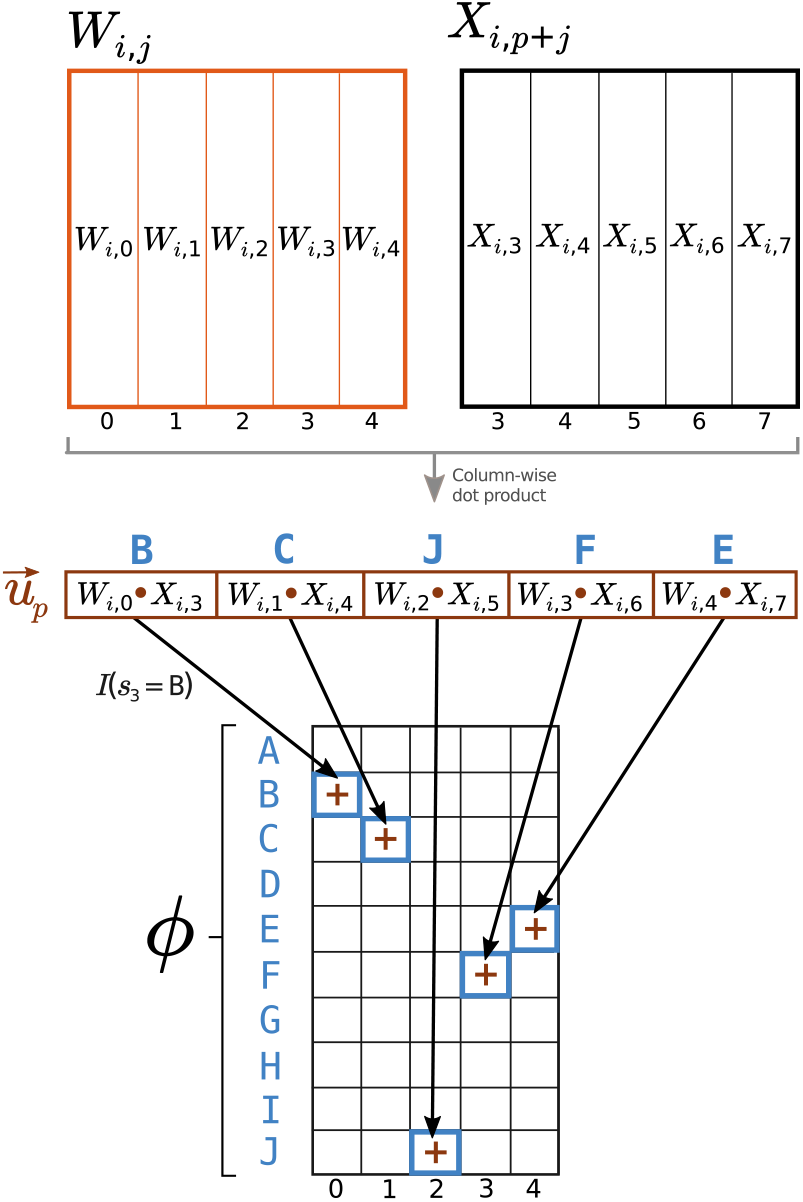
Example computation of *ϕ* for the toy scenario presented in Figure 1. Here, we assume that subsequence BCJFE yielded a maximum convolution value at position *p* = 3. For each overlapping column pair between *W* and *X*, the dot product is calculated and stored in the projection vector 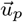 (in brown). Then, elements of 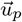 are summed into *ϕ*, a matrix whose coordinates are indexed by (letter, position) pairs, using an indicator function *I* as row mapper (in this figure, an example for B at *s*3 is shown). With this dot product accumulation, the contribution to the projection of *S* onto *ϕ* is leveraged, improving its resolution.

## Materials and Methods

In the context of peptide-MHC interactions, the aforementioned approach can be employed to project binding sequences *s* and identify potential binding motifs that may be distributed across various positions in the input space. However, for this to be effective, it is crucial that the filter weights are precisely adjusted to detect these motifs. If we consider convolutional neural networks as collections of convolutional layers, and each layer as a set of convolutional filters, it becomes feasible to construct a CNN architecture, optimize its filter weights via Gradient Descent, and subsequently compute their projections *ϕ* over a given input peptide space *S*. To begin examining potential experimental scenarios, we adapted the NetPhosPan architecture to the peptide-MHC system (Figure 4), utilizing the Keras deep learning API [Chollet, 2015].

**Figure 4:**
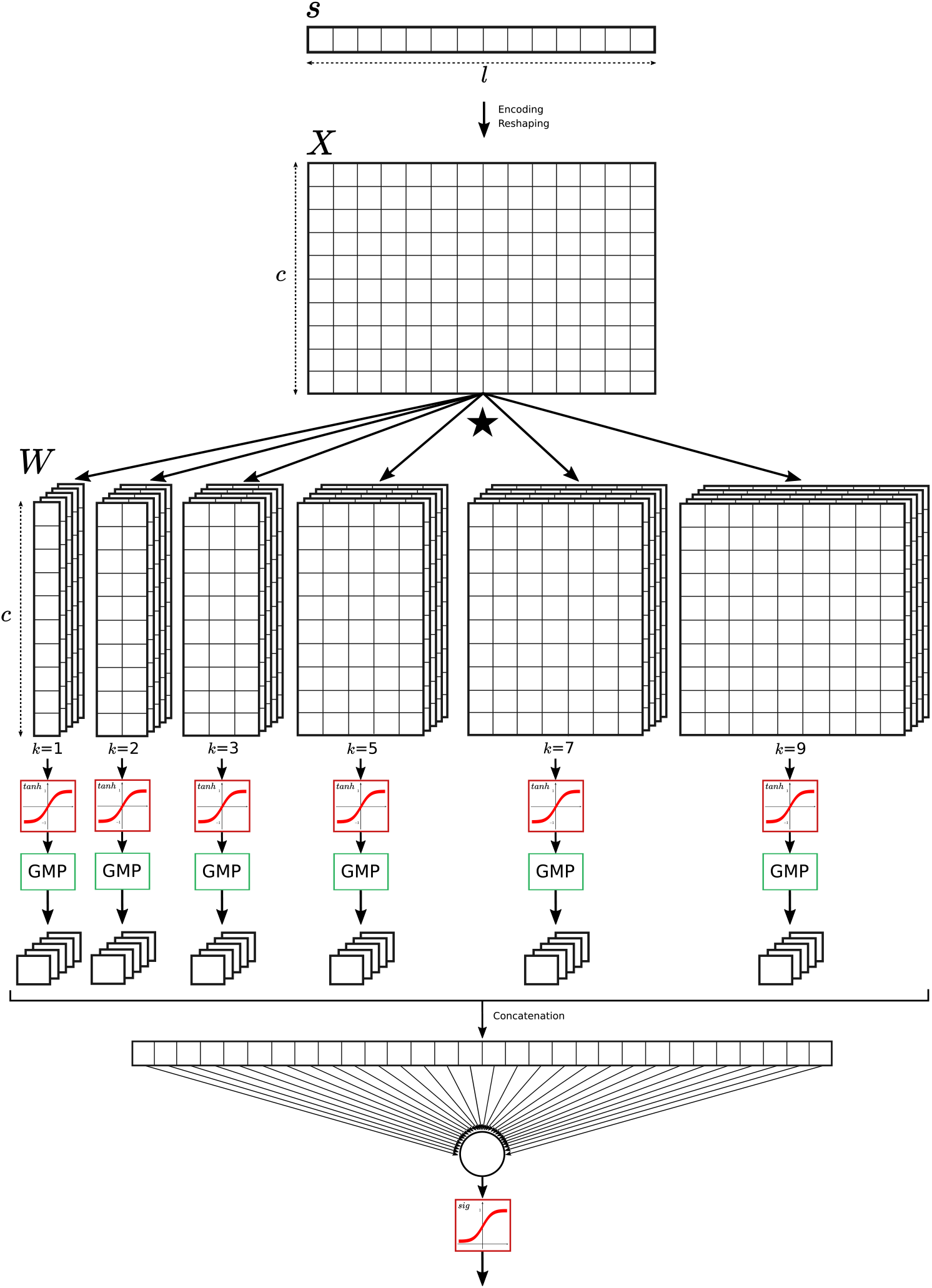
Architectural representation of the proposed model. From top to bottom, an input sequence *s* of length *l* assembled from an alphabet of length *c* is encoded and reshaped into a matrix *X* in order to fit the convolutional layers input shape (*c, l*). For MHC-I and MHC-II, *l* = 14 and *l* = 21, respectively. Also, BLOSUM encoding implies *c* = 20. Six convolutional layers of kernel sizes 1, 2, 3, 5, 7 and 9 (with 5 filters each) are fed the input. CLs outputs are then fed to *tanh* activation functions, and then into `GlobalMaxPool()` (GMP) blocks. The output of each GMP are 5 scalars (each one associated to an upstream filter), which become concatenated into a unique vector of length 30 (six kernels times five filters each) and fed to the output neuron, whose activation function is a sigmoid.

Our approach consists of an allele-specific ANN composed, essentially, of a convolution module with six parallel convolutional layers each of kernel sizes 1, 2, 3, 5, 7, or 9 (in the NetPhosPhan model, kernels of length 3, 5, and 7 were used; we here add a 9-kernel to account for 9-mer binding cores, and 1- and 2-kernels to scan for potentially smaller patterns). The chosen activation function for all CLs was hyperbolic tangent (`tanh`), and the padding was set to valid (this means that, after convolving a peptide of length *l* with a filter of length *k*, the resulting vector will not be zero-padded to the right, and thus will have *l* − *k* + 1 positions). Each activated CL is then fed into a `GlobalMaxPool()` operation, which extracts the maximum value of the activated convolution output. Then, all pooled values become concatenated into a single vector and fed to a single output neuron with sigmoid activation. Since the proposed architecture has multiple convolutional filters *f* ∈ *F* (with *F* being the collection of all filters), we will refer to their projections as *ϕ*_*f*_. Also, since a single output neuron and no hidden layer is present in the proposed model (Figure 4), a weighted projection 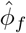 of *S* onto *f* can be computed as:

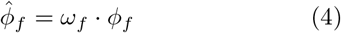

where *ω*_*f*_ is the weight connecting filter *f* (after `GlobalMaxPool()`) to the output neuron (added to account for the network’s assigned importance to). Since in our experimental setup *S* corresponds to a list of specific MHC-restricted peptides, 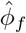 shall display a filter’s “viewpoint” of such MHC binding preferences. Thanks to this, meaningful information regarding receptor motifs might be extracted from such a unique perspective. Also, with the above expression, different 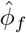 matrices may be combined together to generate composite projections for any filter combination. For instance, the full network projection can be calculated as the sum of all weighted projections 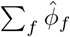.

Having defined the architecture and projection computation procedures, we next proceeded to the training step. A total of six models were trained, accounting for HLA-A*02:01, HLA-A*01:01 and HLA-B*08:01 (HLA-I); and HLA-DRB1*01:01, HLA-DRB1*03:01 and HLA-DRB1*11:04 (HLA-II). Data for the class I system was extracted from the clustering output of NNAlign MA training, whileas data for class II was obtained from the clustering output of NetMHCIIpan-4.0 training. Positive peptides were labeled with a target value of 1, while negative peptides with a target value of 0. Negative enrichment was 5x the quantity of the most abundant positive length, for all lengths. Peptides were padded to a maximum length of 14 (in the case of MHC-I), and a maximum length of 21 (in the case of MHC-II). The chosen peptide encoding was BLOSUM50, rescaled by a factor of 5; the padding character “X” (wildcard amino acid) was encoded with a vector of length 20 and values of −1. Optimization was done with Batch Gradient Descent, with batch size of 64, learning rate of 0.05 and binary cross-entropy as loss function. Training was conducted for 200 epochs with an early stopping of 40 epochs, following a 5-fold cross validation schema with sequence homology reduction. A total of 5 filters per convolutional kernel were used for all models, except HLA-DRB1*11:04, for which we used 10.

## Results

All six trained models exhibited strong performance, achieving an average cross-validated ROC AUC of 0.941 and a PRC AUC of 0.793 (see Supplementary Figure 1 for further details). The cross-validated metrics for HLA-B08:01 are specifically illustrated in Figure 5. With successful training confirmed, the next objective was to determine optimal projections for all models and reconstruct their corresponding MHC binding motifs. Using HLA-B08:01 as an illustrative example, such a projection should closely resemble the established HLA-B08:01 binding motif (Figure 6), which will serve as our target PSSM.

**Figure 5:**
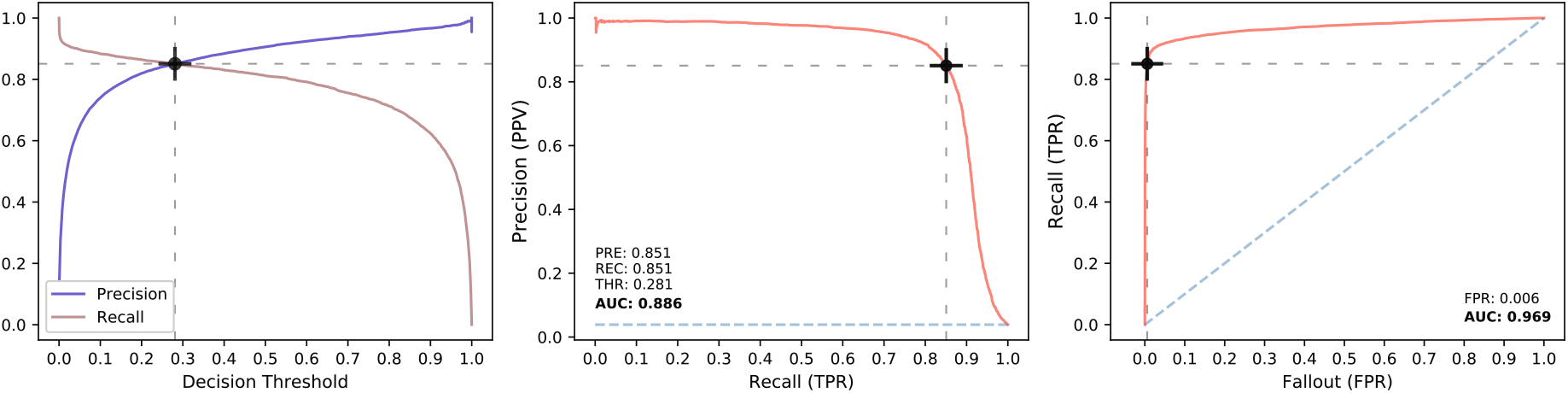
Cross-validation metrics for the HLA-B*08:01 model. From left to right: Precision and Recall curves as a function of the decision threshold, Precision-Recall curve and ROC curve. The black cross indicates the chosen operation point for the model (in this case, the intersection of the precision and recall curves). The AUC values for PRC and ROC are shown in bold.

**Figure 6:**
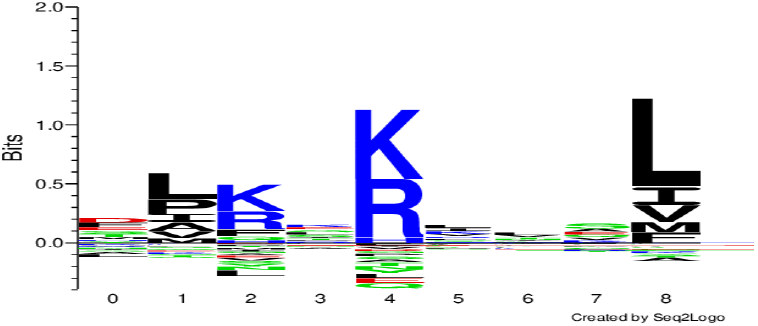
HLA-B*08:01 binding motif. Known logo, extracted from the top 0.1% of 200.000 random peptide predictions.

To do this, four cumulative projection alternatives were computed for all filters (Figure 7). The first panel shows that omitting *ω*_*f*_ results in a divergent pattern with little similarity to the target logo (Figure 6), featuring a monotonically decreasing information content shape. Including the absolute value of *ω*_*f*_ (second panel) reduces this monotony, allowing the P5 anchor to emerge. Using the sign of *ω*_*f*_ as weight (third panel) produces a sharper P5, introduces P3, and accurately positions the P9 anchor. On the other hand, employing *ω*_*f*_ (fourth panel) yields the best result of all, with overall more informative and crisp anchors, but also with correct, depleted enrichments at unimportant positions as well. Refer to Supplementary Figure 2 for similar results for the remaining models.

**Figure 7:**
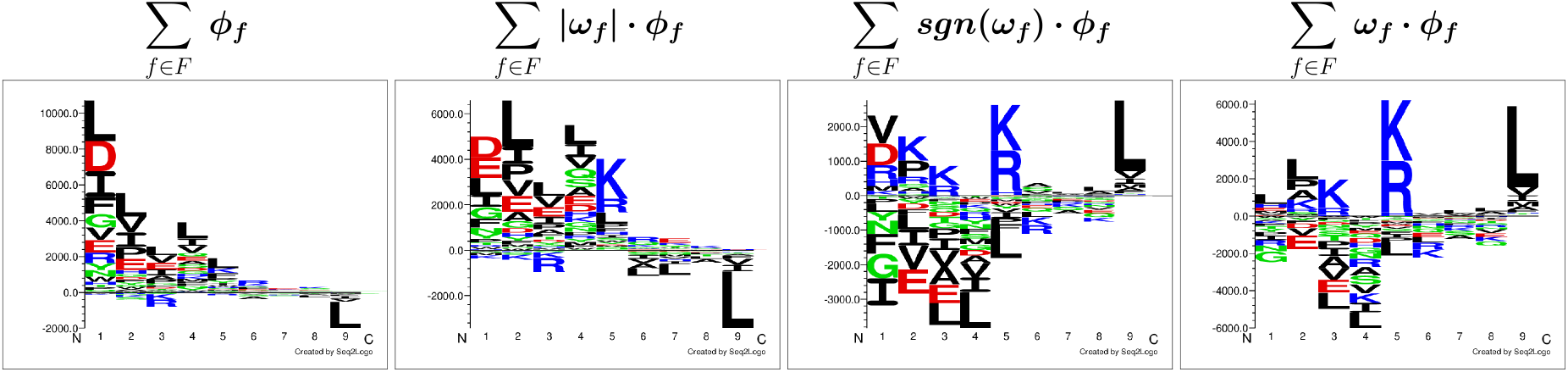
Four different cumulative projections for the model trained on HLA-B*08:01 data. From left to right: no weighting, weighting using the absolute value of the output neuron weight, weighting using the sign of the output neuron weight, and weighting using the output neuron weight.

These results underscore the importance of a filter’s associated output weight in determining the internal behavior of the architecture. If this were not the case, the summed projections displayed in Figure 7 would show similar characteristics across all four weighting scenarios. This observation suggests that, during training, the network assigns varying levels of importance to each filter by adjusting the magnitude of its weight, such that filters with larger weights are deemed more significant, while those with smaller weights are less so. In addition to magnitude, the sign of the weight is also crucial. This can be understood by considering the outputs of the hyperbolic tangent activation function, which range from [−1, 1]. The network uses this range to separate positive and negative classes by pushing positive predictions toward 1 and negative ones toward −1, or vice versa. This approach enables the network to assign different classes to distinct regions in its internal representation, facilitating effective separation. To align with the actual MHC binding motif preferences, the learning process enforces either a positive or negative sign on each weight, thereby flipping the filter outputs accordingly when they are summed by the output neuron (refer to Figure 8 for an example).

**Figure 8:**
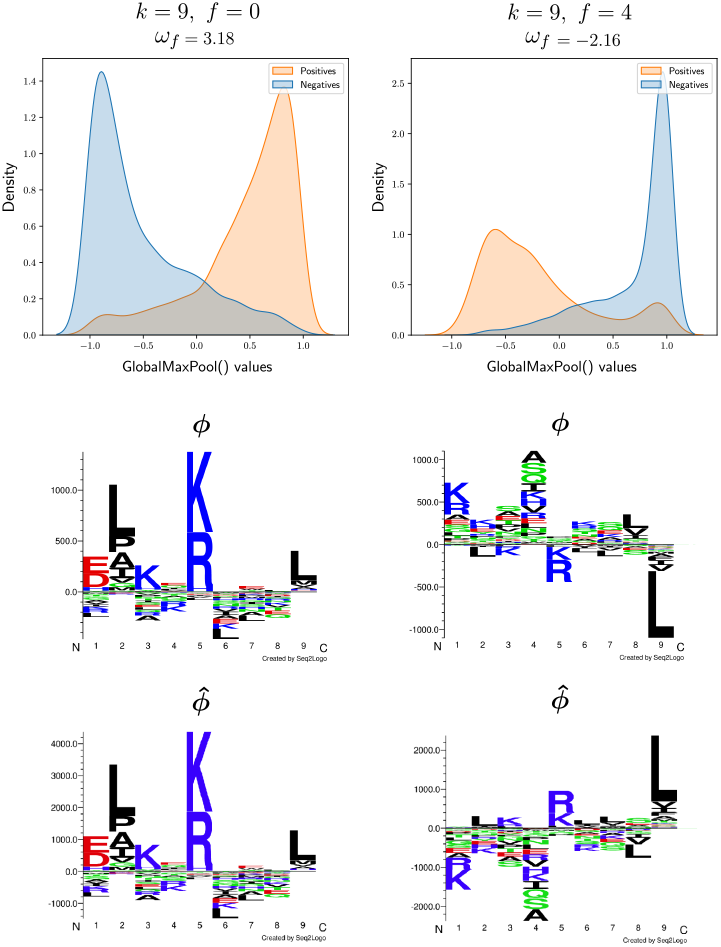
Role of the output weight *ω*_*f*_ in generating consistent internal projections. On the top row, density estimations for `GlobalMaxPool()` outputs are shown for the first filter (*f* = 0, first column) and last filter (*f* = 4, second column) of the convolutional layer of kernel size *k* = 9 (the corresponding output weight is shown on top of the plots). Densities corresponding to the positive and negative training sets are plotted separately. Given that GMP receives the output of a `tanh` activation function as input, such values will be bound between −1 and 1, guiding the network to squish class separation on these extremes. Notice how such separation can be obtained disregarding the tanh sign: for *f* = 0, positives are squished towards 1, whereas for *f* = 4, towards −1. As a direct consequence of this, projections *ϕ* for both filters (middle row) will be x-axis mirrored (both will display similar anchors, but on opposite sides). When applying the weighting 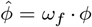 (bottom row), the mirroring becomes fixed, and now important anchors point towards the same direction (amplitudes become corrected too). With this, the summation of 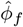 over all *f* ∈ *F* will yield a more sound PSSM, as seen on the rightmost logo of Figure 7. Note: density plots display values outside the [−1,1] interval. This is an artifact of the visualization; these do not exist in reality.

Reiterating the importance of *w*_*f*_, Figure 9 presents a side-by-side comparison between the logo derived from the raw input data, the sum of all 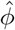, and the logo obtained from the top predictions. Here, a strong resemblance can be observed between our cumulative weighted projection and the NetMHCpan-4.1 logo. In contrast, the logo generated from the raw peptide data exhibits more divergent characteristics compared to the projection, suggesting that this method indeed captures meaningful and valid information. Additionally, it is important to note that, unlike Equation 4, any 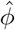 (and particularly the cumulative blend 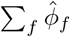) represents a pseudo-PSSM that encodes the network’s internal representation of specific amino acid combinations, which cannot be interpreted as classical information content. In practical terms, this implies that the y-axis of the logo representation of any projection combination will show dimensionless values, unlike the NetMHCpan logos, which represent information units. Therefore, the logo comparisons in this study are made qualitatively through visual inspection, rather than using objective quantification methods such as Spearman’s rank correlation.

**Figure 9:**
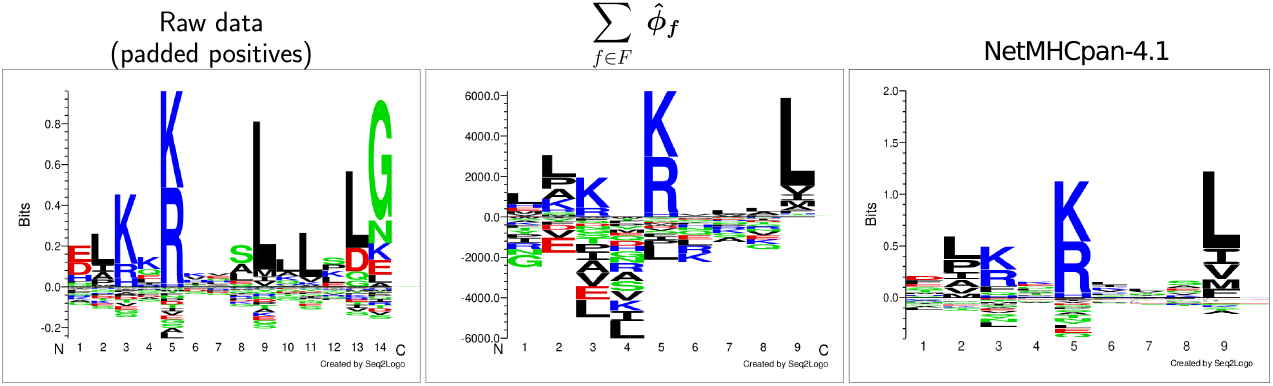
HLA-B*08:01 logo comparison between raw input data (left), summation of all weighted projections (middle) and NetMHCpan-4.1 top 0.1% predictions (right). Raw data logo has 14 positions in total because all input peptides are padded to such length.

For a second example, we repeated the projection extraction process using the model trained on HLA-A*01:01 binding data and generated the corresponding logo (Figure 10). This scenario presented notable differences compared to previous results. Most significantly, the raw data logo for this allele showed a pronounced Tyrosine enrichment towards the C-terminus, which is a direct consequence of the padding strategy used, keeping the N-terminus fixed and causing P9 anchors across input peptides to become misaligned. This misalignment unfortunately affects the cumulative projection, leading to significant discrepancies at P9 compared to the NetMHCpan logo. To address this misalignment issue, we developed a simple alignment pipeline. First, we selected the projection with the highest amplitude (or with the greatest “pseudo-information” content) from the convolutional layer with the longest kernel size to serve as the alignment template, 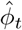.

**Figure 10:**
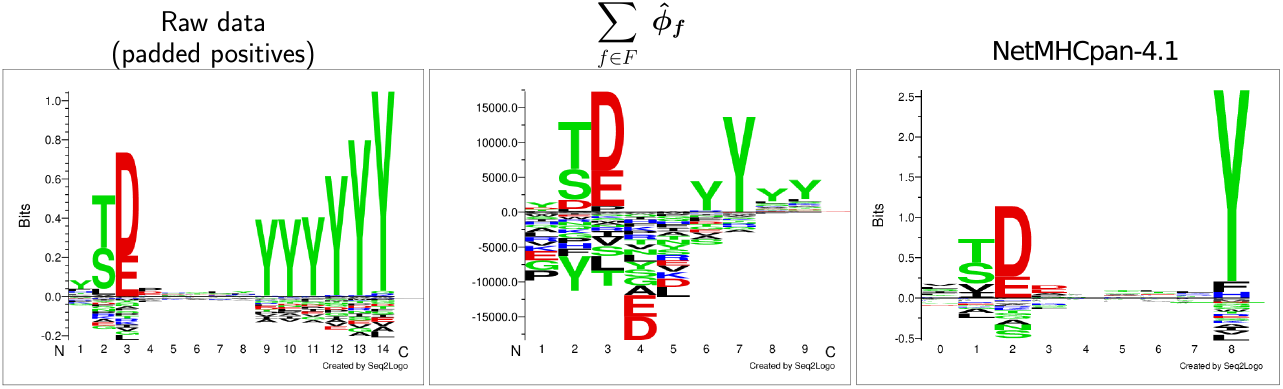
HLA-A*01:01 logo comparison between raw input data (left), summation of all weighted projections (middle) and NetMHCpan-4.1 top 0.1% predictions (right). Raw data logo has 14 positions in total because all input peptides are padded to such length.

Next, all remaining projections 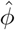 were sorted from highest to lowest based on their maximum positive amplitude within each kernel size group *k*. Starting with the highest value of *k*, a 2-dimensional cross-correlation was calculated between the alignment template 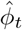and the first 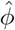 from the sorted list, resulting in the extraction of an offset *o* that maximizes the correlation. This offset *o* is then used to apply a correction to 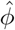, aligning it with the template. The offset-corrected 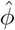 is sub-sequently added to 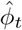, updating the template. This correlation and correction procedure is repeated for the next projection in the sorted list, and the process continues until all projections are consumed. Ultimately, 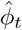 becomes the final aligned version. The results of applying this alignment pipeline to all trained models are presented in Figure 11.

**Figure 11:**
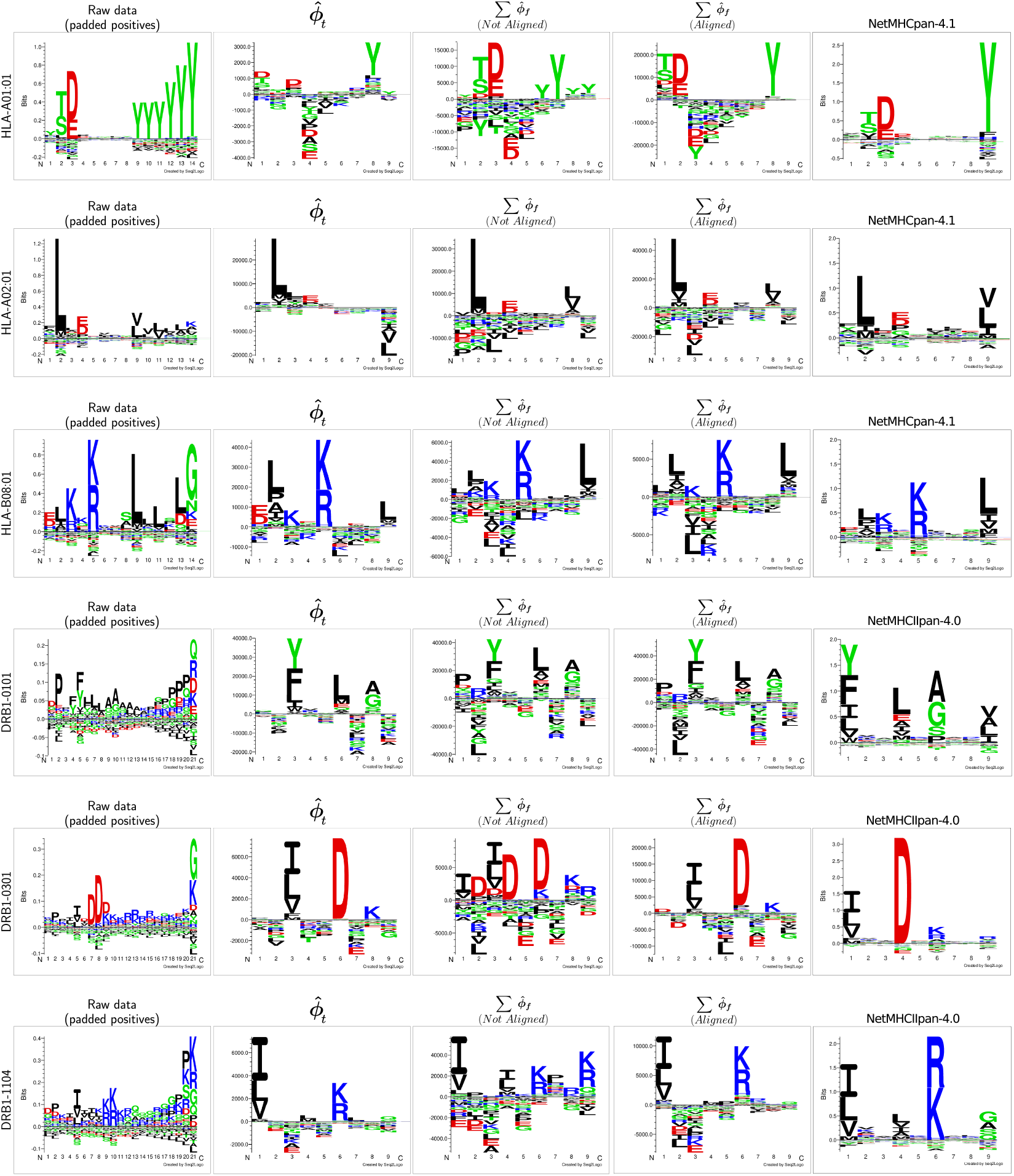
Comparison between logos for raw input data (first column), alignment template (second column), weighted projection accumulation (third column), aligned weighted projection accumulation (fourth column), and known NetMHCpan logo (fifth column), for all employed models (one per row).

As shown in Figure 11, the final outcome of projecting *S* onto the filter space of the trained models varies significantly depending on the HLA molecule being analyzed. The most straightforward case was HLA-B*08:01, whose projection appeared quite accurate even before alignment, with only minor discrepancies such as K/R enrichment at P2 or a slight P9 anchor appearing at P7 and P8. After alignment, these minor errors were resolved, although R at P3 was lost in the process. For HLA-A01:01, noticeable misalignments were present in the initial solution, evident from the fragmented “ghost anchors” at P9, which became unified after alignment. Interestingly, the sum of the aligned projections begins at P2 of the NetMHCpan logo, but the positional relationships are still well preserved. The scenario for HLA-A02:01 was different, as alignment did not result in substantial changes apart from an increase in amplitude, and the misplaced P9 anchor could not be corrected.

In the case of the MHC-II system, both pre- and post-alignment projections for HLA-DRB1*01:01 were quite similar, showing no significant qualitative improvement. However, both projections started two positions to the left of P1 in the NetMHCIIpan logo, resulting in the final motif being two positions shorter toward the C-terminus.

We believe this shift may be due to a consistent Proline (P) signal upstream of P1, which aligns with previously characterized MHC-II antigen processing signals [Barra et al., 2018]. For HLA-DRB103:01, the alignment process provided a clear benefit by unifying the scattered P1, P4, and P6 anchors. As with HLA-DRB101:01, the two-position shift was also observed here. The final allele, HLA-DRB111:04, also benefited from alignment, which produced well-defined P1 and P6 anchors. However, less prominent anchors, such as P4 and P9, appeared diluted, being present but with low magnitudes in the aligned 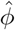. Notably, unlike HLA-DRB103:01 and HLA-DRB1*11:04, we did not observe the two-position shift, and all anchors remained well-aligned with the NetMHCIIpan logo.

In conclusion, regardless of individual model gains, losses, or limitations, the aligned 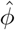 consistently represents a more accurate binding motif estimation compared to the raw input data. For a more detailed overview of how alignments were generated by summing offset-corrected projections, refer to Supplementary Figures 3, 4, 5, 6, 7, and 8.

## Discussion

With the results presented above, it becomes clear that neural networks develop complex internal representations of the input data they are given. Due to the sparse and intricate nature of these abstraction spaces, where information is embedded within weights and connections, understanding the internal language of neural networks is far from straight-forward. In this work, we have made a modest attempt to gain insight into these processes, focusing specifically on the case of 1-dimensional convolutional neural networks.

Since this deep learning approach is particularly well-suited for analyzing peptide-MHC binding interactions, we constructed a basic architecture to investigate the problem in a controlled environment. We leveraged the convolution operator to project the input set onto the weight space of the convolutional filters, enabling us to visualize the internal constructs of the network using sequence logo representations. We trained a total of six different peptide-MHC binding prediction models. The results were promising, as the computed projections showed considerable resemblance to known experimental binding motifs. Nevertheless, we had to introduce an offset correction step to address a mis-alignment issue in some anchors, which ultimately improved the alignment for HLA-II, a more complex system.

Based on these findings, we believe that this novel way of examining 1D convolutions could expand our understanding of how artificial neural networks build, integrate, and utilize internal representations of input data. The work presented here represents an initial exploration, leaving considerable room for further improvements and scaling. As seen in Supplementary Figures 3, 4, 5, 6, 7, and 8, many individual projections do not seem to convey informative content (showing no clear patterns or having low amplitude), suggesting that not all filters are optimized equally. This opens up questions about whether this behavior reflects a genuine issue or merely the normal functioning of the network. Future work could explore this using regularization techniques, such as L1/L2 penalties or dropout layers, and measure how these impact the projection shapes.

Furthermore, since output weights are central to this approach and are influenced by filter activation calculations, it is crucial to examine the role of different activation functions in generating projections. In our study, we used the hyperbolic tangent (tanh), which restricts outputs to the range [−1, 1], necessitating both positive and negative weight adjustments to regulate filter responses. Exploring alternative activation functions such as ReLU, Leaky ReLU, or Swish could affect the way networks represent information, potentially leading to richer and more diverse internal representations. ReLU, for instance, could reduce inactive filters and emphasize more relevant features, while Swish could offer greater flexibility with both positive and negative outputs, potentially capturing more nuanced biological motifs.

Expanding the current architecture to a pan-specific framework is also an important direction for future work. In such a setup, a single convolutional filter would need to capture information from multiple MHC molecules, thus losing filter-MHC exclusivity and potentially leading to new abstraction patterns. However, directly adapting the current architecture to train a pan-specific model may present performance issues, primarily because the existing setup uses a single output neuron, which might be insufficient for processing multiple MHC molecules. Expanding the architecture to include multiple output neurons, or even additional hidden layers, would likely enhance predictive performance. It is important to consider, though, that by doing so the weighting schema for 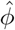 would need to be adapted, as it currently assumes a single output neuron.

In conclusion, while the landscape of these deep learning approaches is broad and complex, it shows immense promise and is well worth further exploration. The use of convolutional neural networks to model peptide-MHC interactions has already shown effectiveness in capturing complex biological patterns essential for understanding immune responses. By refining model architectures and training schemas, there is significant potential to improve predictions and biological insight. The use of advanced activation functions, regularization, and multi-resolution methods offers exciting avenues to address current limitations, such as identifying longer motifs or enhancing interpretability. Ultimately, by optimizing these models and ensuring their transparency, we can enhance both their predictive power and their applicability in clinical settings, contributing to a deeper understanding of immunology and advancing healthcare solutions.

