## supplementary_figures for "Deep Learning-Based Motif Discovery in Major Histocompatibility Complex: A Primer"

### Supplementary Material

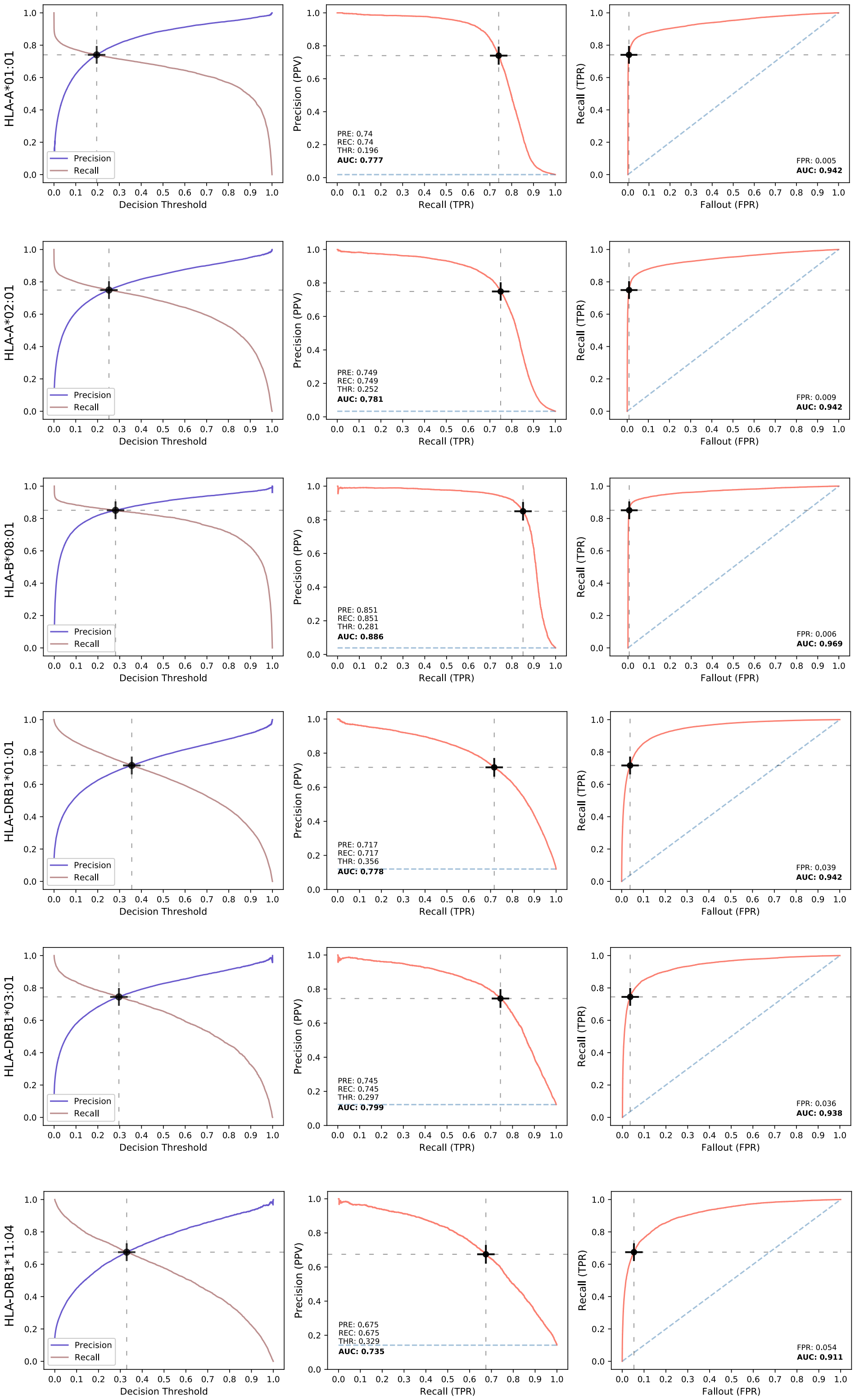


Supplementary Figure 1: **Cross-validation metrics for all trained models (one per row).** From left to right columns: Precision and Recall curves as a function of the decision threshold, Precision-Recall curve and ROC curve. The black cross indicates the chosen operation point for the model (in this case, the intersection of the precision and recall curves). The AUC values for PRC and ROC are shown in bold.

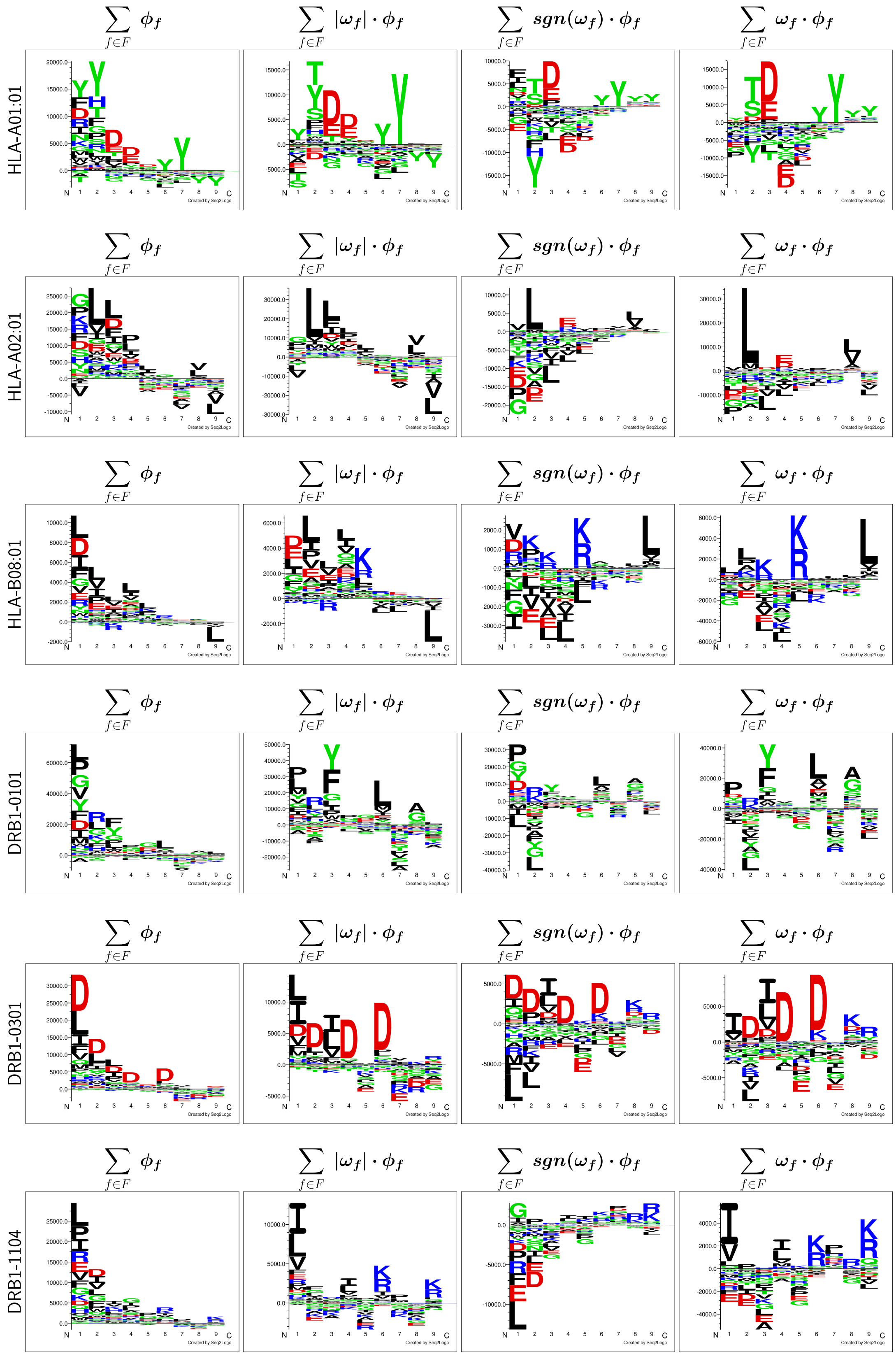


Supplementary Figure 2: **Logos resulting from different $\phi_f$ weighting approaches, for each one of the trained models (indexed by rows).** From left to right columns: no weighting, weighting using the absolute value of the output neuron weight, weighting using the sign of the output neuron weight, and weighting using the output neuron weight.

### HLA-A01:01

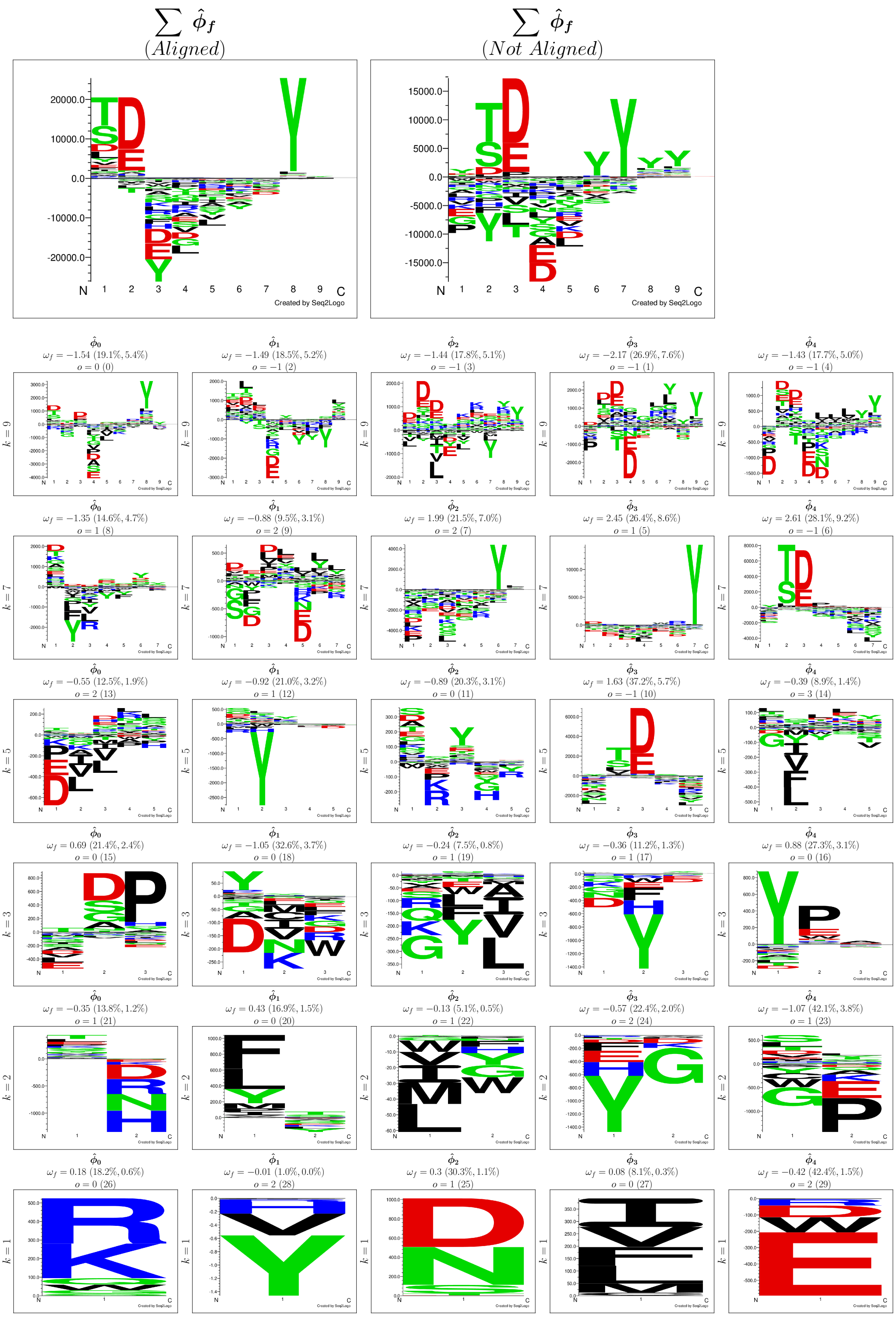


Supplementary Figure 3: **Full projection panel for the HLA-A*01:01 molecule.** On top, the aligned and not aligned accumulated weighted projections are shown in the left and in the right, respectively. Below, all the individual weighted projections $\hat{\phi}$ for all filters (indexed by columns) in each convolutional layer of kernel size $k$ (indexed by rows) are shown. On top of each projection, the value of $\omega_f$ (weight connecting the corresponding filter to the output neuron) is shown. Next to this, and in between parentheses, the contribution percentage of $\omega_f$ to the convolutional layer and to the full network are displayed. Below, the offset correction value $o$ is exhibited; next to it, in between parentheses, the order in which the projection was added to the cumulative projection is shown.

### HLA-A02:01

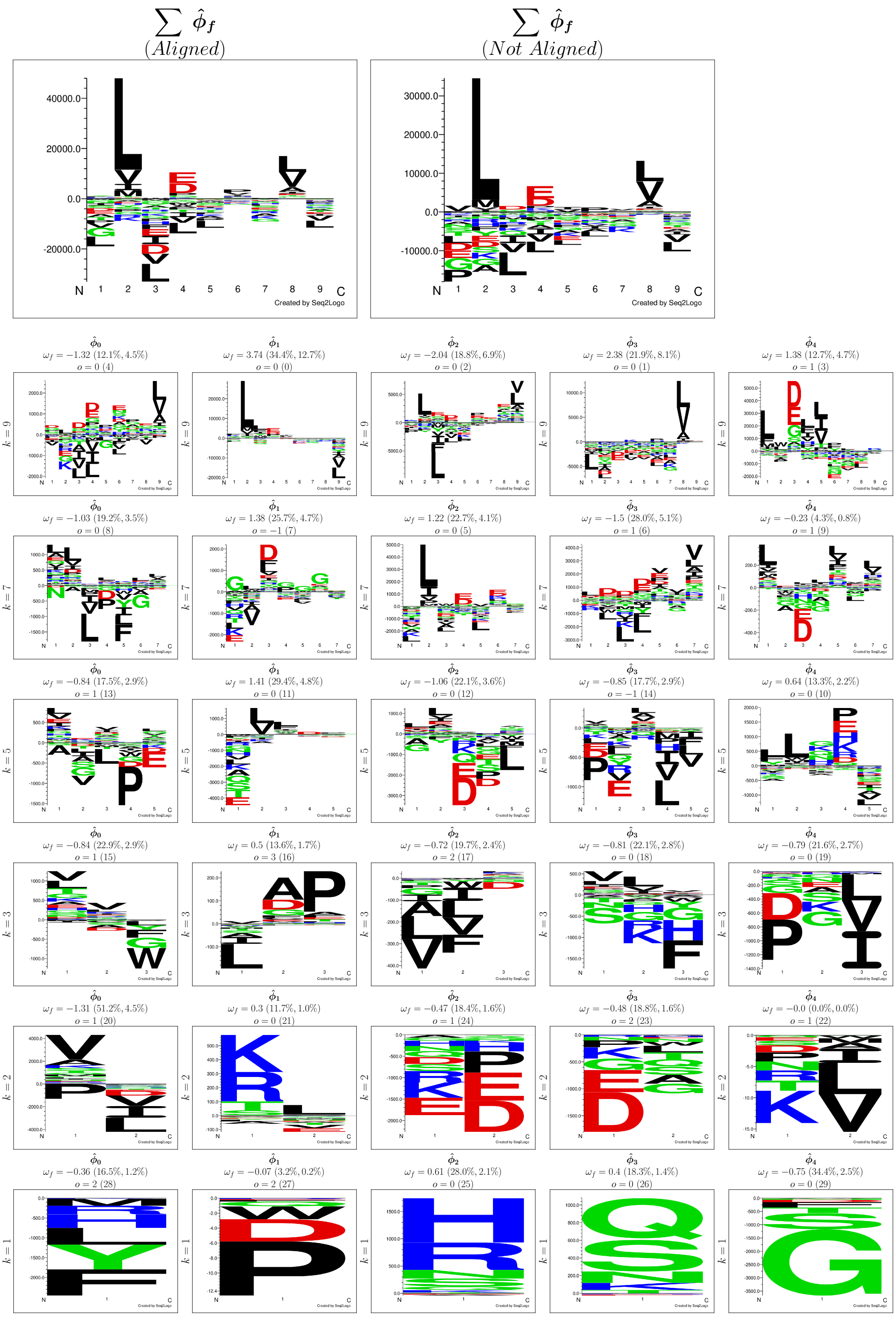


Supplementary Figure 4: **Full projection panel for the HLA-A*02:01 molecule.** On top, the aligned and not aligned accumulated weighted projections are shown in the left and in the right, respectively. Below, all the individual weighted projections $\hat{\phi}$ for all filters (indexed by columns) in each convolutional layer of kernel size $k$ (indexed by rows) are shown. On top of each projection, the value of $\omega_f$ (weight connecting the corresponding filter to the output neuron) is shown. Next to this, and in between parentheses, the contribution percentage of $\omega_f$ to the convolutional layer and to the full network are displayed. Below, the offset correction value $o$ is exhibited; next to it, in between parentheses, the order in which the projection was added to the cumulative projection is shown.

### HLA-B08:01

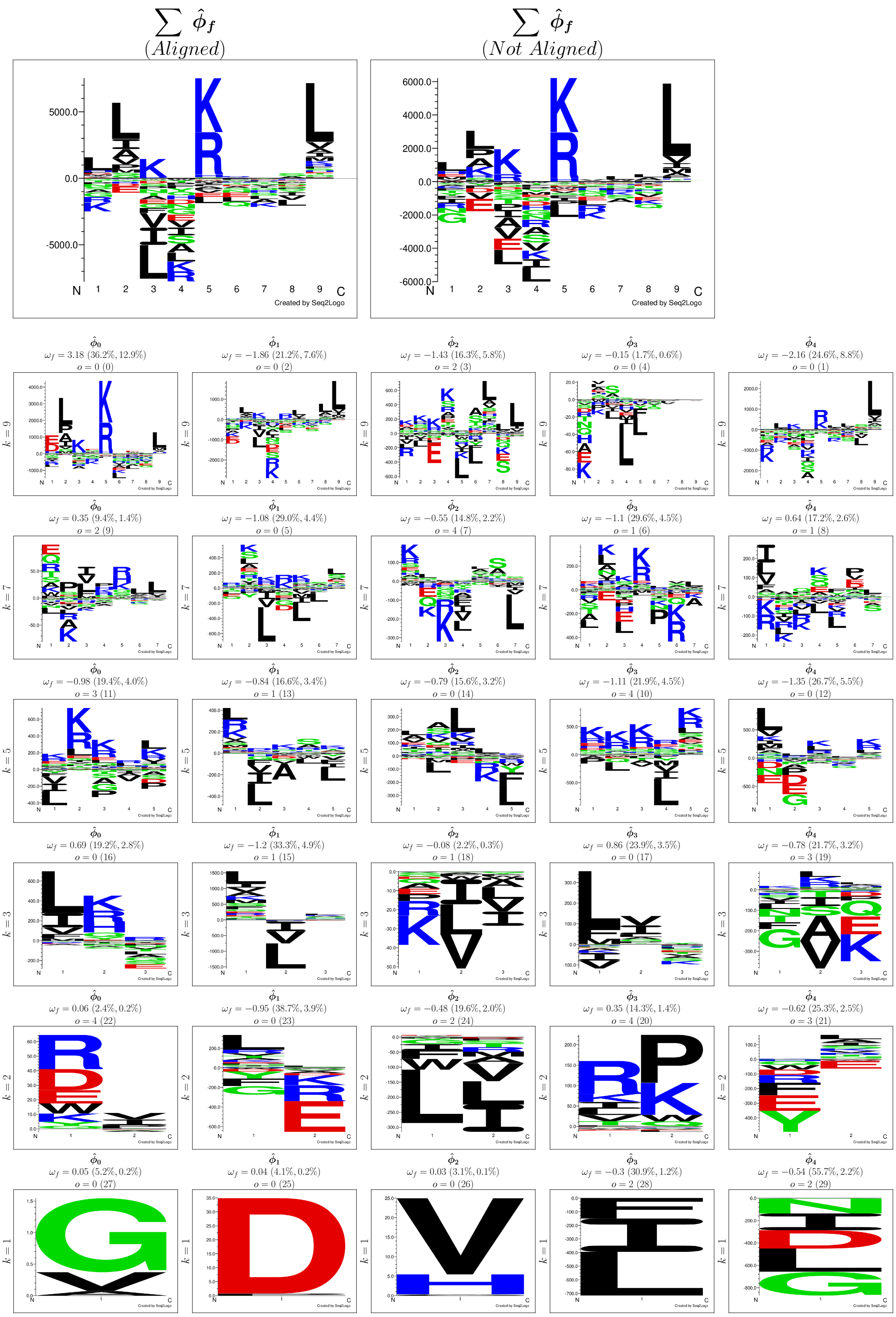


Supplementary Figure 5: **Full projection panel for the HLA-B*08:01 molecule.** On top, the aligned and not aligned accumulated weighted projections are shown in the left and in the right, respectively. Below, all the individual weighted projections $\hat{\phi}$ for all filters (indexed by columns) in each convolutional layer of kernel size $k$ (indexed by rows) are shown. On top of each projection, the value of $\omega_f$ (weight connecting the corresponding filter to the output neuron) is shown. Next to this, and in between parentheses, the contribution percentage of $\omega_f$ to the convolutional layer and to the full network are displayed. Below, the offset correction value $o$ is exhibited; next to it, in between parentheses, the order in which the projection was added to the cumulative projection is shown.

### DRB1-0101

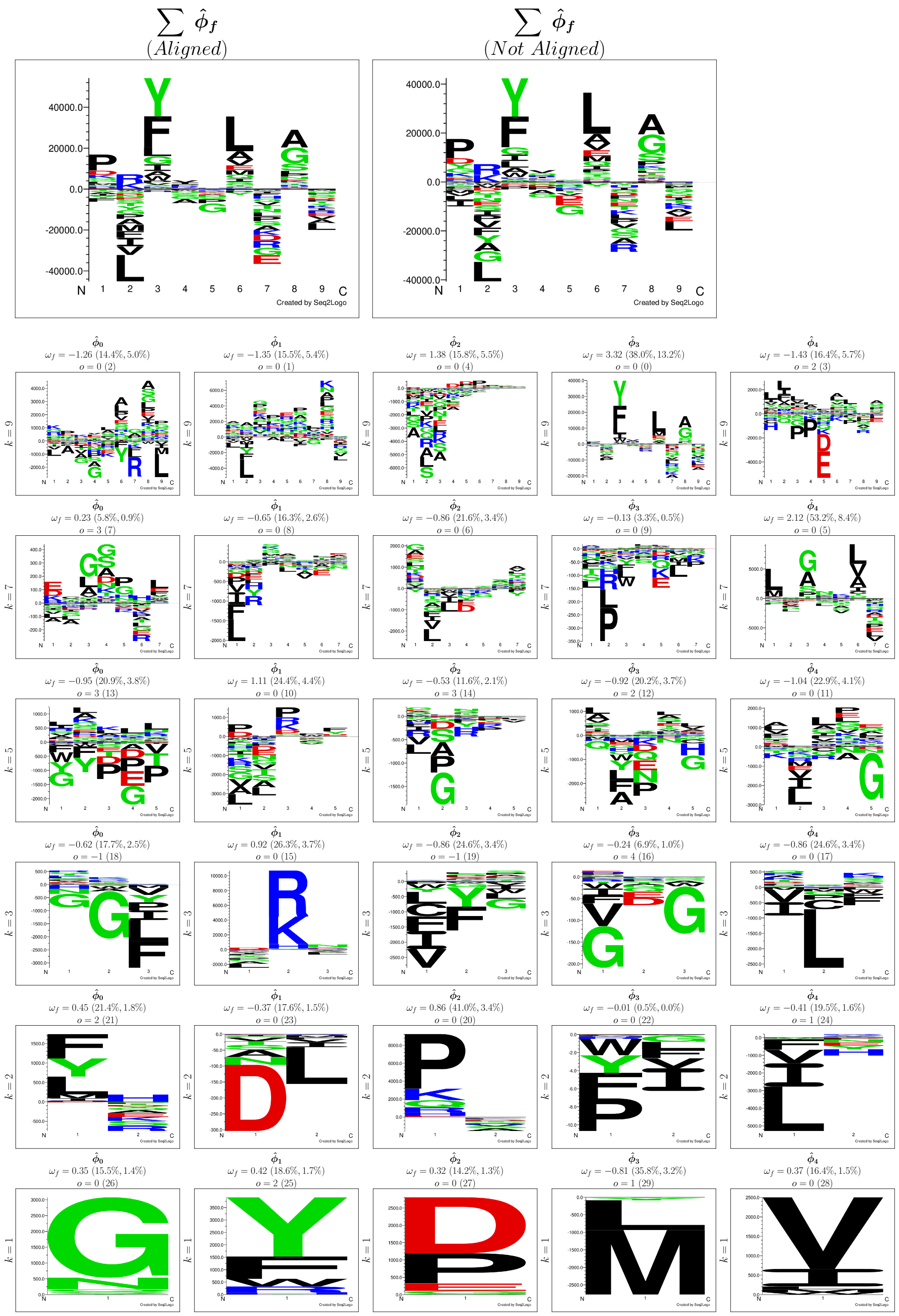


Supplementary Figure 6: **Full projection panel for the HLA-DRB1*01:01 molecule.** On top, the aligned and not aligned accumulated weighted projections are shown in the left and in the right, respectively. Below, all the individual weighted projections $\hat{\phi}$ for all filters (indexed by columns) in each convolutional layer of kernel size $k$ (indexed by rows) are shown. On top of each projection, the value of $\omega_f$ (weight connecting the corresponding filter to the output neuron) is shown. Next to this, and in between parentheses, the contribution percentage of $\omega_f$ to the convolutional layer and to the full network are displayed. Below, the offset correction value $o$ is exhibited; next to it, in between parentheses, the order in which the projection was added to the cumulative projection is shown.

### DRB1-0301

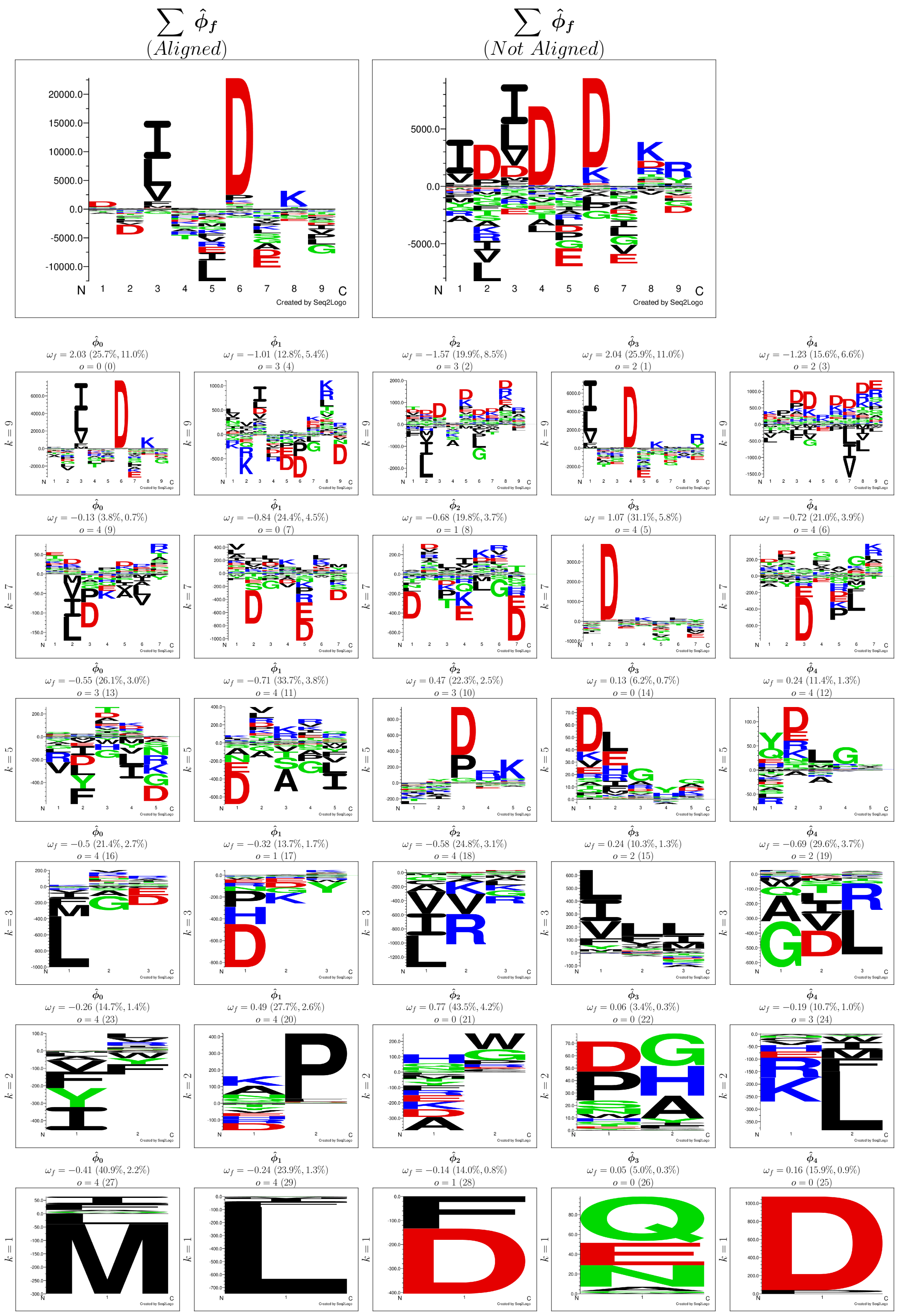


Supplementary Figure 7: **Full projection panel for the HLA-DRB1*03:01 molecule.** On top, the aligned and not aligned accumulated weighted projections are shown in the left and in the right, respectively. Below, all the individual weighted projections $\hat{\phi}$ for all filters (indexed by columns) in each convolutional layer of kernel size $k$ (indexed by rows) are shown. On top of each projection, the value of $\omega_f$ (weight connecting the corresponding filter to the output neuron) is shown. Next to this, and in between parentheses, the contribution percentage of $\omega_f$ to the convolutional layer and to the full network are displayed. Below, the offset correction value $o$ is exhibited; next to it, in between parentheses, the order in which the projection was added to the cumulative projection is shown.

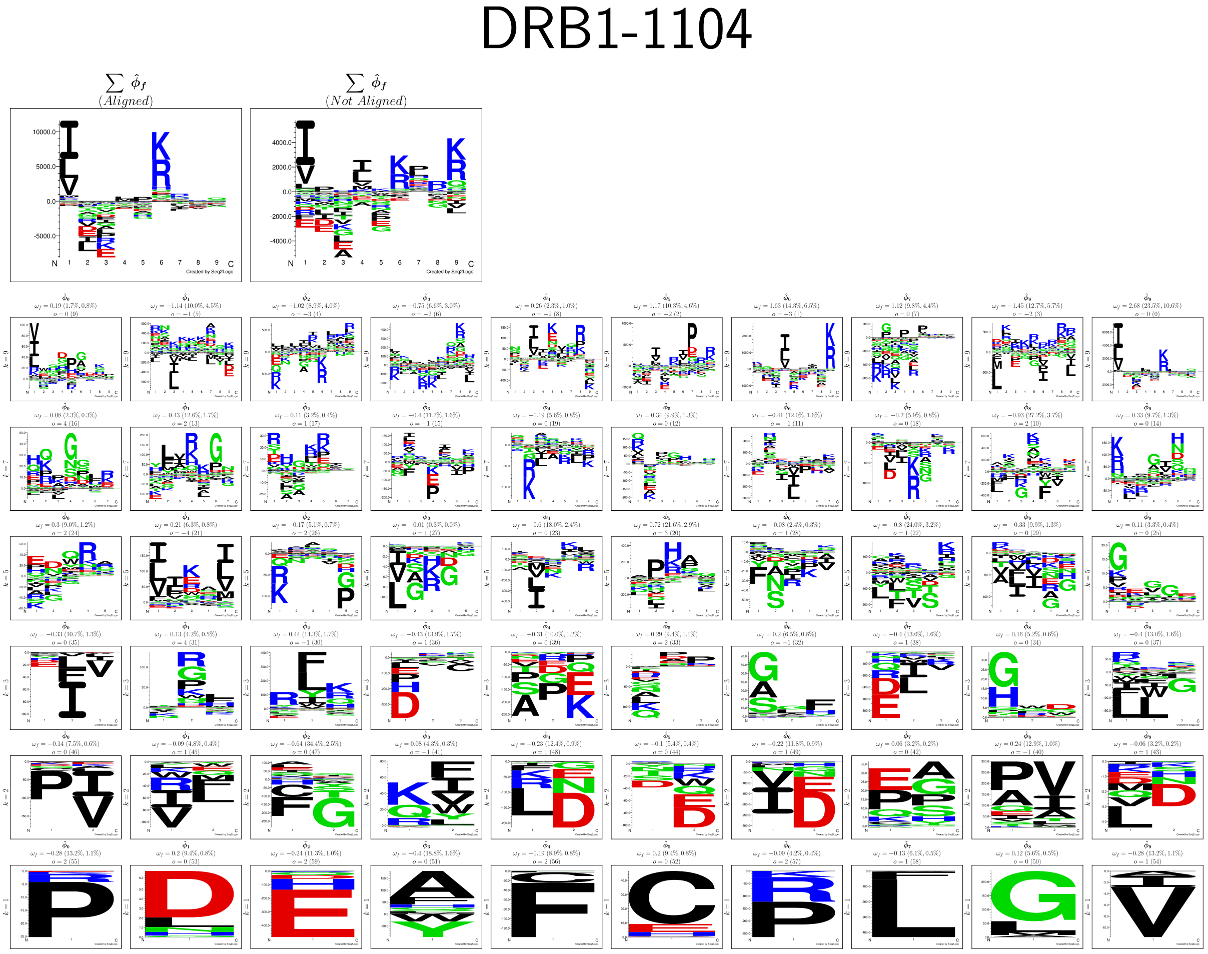

Supplementary Figure 8: **Full projection panel for the HLA-DRB1*11:04 molecule.** On top, the aligned and not aligned accumulated weighted projections are shown in the left and in the right, respectively. Below, all the individual weighted projections $\hat{\phi}$ for all filters (indexed by columns) in each convolutional layer of kernel size $k$ (indexed by rows) are shown. On top of each projection, the value of $\omega_f$ (weight connecting the corresponding filter to the output neuron) is shown. Next to this, and in between parentheses, the contribution percentage of $\omega_f$ to the convolutional layer and to the full network are displayed. Below, the offset correction value $o$ is exhibited; next to it, in between parentheses, the order in which the projection was added to the cumulative projection is shown.
